# Retrograde transport of Akt by a neuronal Rab5-APPL1 endosome

**DOI:** 10.1101/499004

**Authors:** Livia Goto-Silva, Marisa P. McShane, Sara Salinas, Yannis Kalaidzidis, Giampietro Schiavo, Marino Zerial

## Abstract

Long-distance axonal trafficking plays a critical role in neuronal function, and transport defects have been linked to neurodegenerative disorders. Various lines of evidence suggest that the small GTPase Rab5 plays a role in neuronal signaling via early endosomal transport. Here, we characterized the motility of Rab5 endosomes in primary cultures of mouse hippocampal pyramidal cells by live-cell imaging and showed that they exhibit bi-directional long-range motility in axons, with a strong bias toward retrograde transport. Characterization of key Rab5 effectors revealed that endogenous Rabankyrin-5, Rabenosyn-5 and APPL1 are all present in axons. Further analysis of APPL1-positive endosomes showed that, similar to Rab5-endosomes, they display more frequent long-range retrograde than anterograde movement, with the endosomal levels of APPL1 correlated with faster retrograde movement. Interestingly, APPL1-endosomes transport the neurotrophin receptor TrkB and mediate retrograde axonal transport of the kinase Akt1. FRET analysis revealed that APPL1 and Akt1 interact in an endocytosis-dependent manner. We conclude that Rab5-APPL1 endosomes exhibit the hallmarks of axonal signaling endosomes to transport Akt1 in hippocampal pyramidal cells.

## Introduction

The ability of axons to transport cargo over long distances is critical for processes ranging from axon path finding and target innervation, to neuronal survival. Those processes are regulated by the retrograde transport of neurotrophins such as nerve growth factor (NGF) and brain-derived neurotrophic factor (BDNF) ^1^. Mutations or alterations in the expression of genes that enable axonal transport are linked to neurodegenerative diseases, including Charcot–Marie–Tooth disease type 2, Huntington’s disease (HD), amyotrophic lateral sclerosis (ALS) and Alzheimer’s disease (AD) ^2,3^. Abnormal axonal traffic in cortical and hippocampal neurons is an early feature of both AD and HD, and a trigger for synaptic loss, neuronal death and the resultant loss of cognitive abilities ^4–6^. Thus, understanding the mechanisms that control axonal transport of survival and differentiation signals could provide important insights into the physiopathology of the brain.

Fast axonal transport of signaling molecules is largely mediated by endosomes. Endosomes facilitate signaling through rapid retrograde transfer of proteins such as neurotrophins, activated receptor complexes, adaptors and kinases along microtubule tracks ^7–9^. The identity of such endosomal signaling compartments has long been debated. In hippocampal neurons, multi-vesicular bodies transport endocytic cargo retrogradely in the axon ^10^. To determine the molecular identity and signaling properties of endosomes in central nervous system (CNS) neurons, it is necessary to characterize their molecular machinery ^11^. Different studies have provided evidence in favor of both early and late endosomal compartments to transport signaling molecules from the pre-synaptic terminal retrogradely to the cell body ^12^. Key regulators of organelle tethering, docking, signaling, fusion and motility are Rab GTPases and their effectors ^13,14^. Rab5 localizes to early endosomes, which sort cargo to the recycling pathway via Rab4 and Rab11, or to the degradative pathway through Rab7-positive endosomes ^14^. In neurons, both Rab5 and Rab7 were shown to be important for retrograde trafficking. In dorsal root ganglion neurons, retrograde movement of NGF has been associated with Rab5-positive endosomes ^9^. In contrast, Rab7-positive endosomes retrogradely transport GFP-BDNF in motor neurons^12^.

However, the identity of endosomal compartments is best defined by the combination of Rab GTPases with their specific effectors, as this can differ significantly between organelles in space and time. For example, in non-polarized cells two distinct populations of Rab5-positive early endosomes coexist and dynamically exchange cargo over time, one containing the canonical Rab5 effector EEA1 and the other harboring APPL1 ^15,16^. In hepatocytes, EEA1- and APPL1-positive endosomes have distinct distributions that depend on the organization of the actin cytoskeleton and the nutritional state ^17^.

EEA1, which functions as a tethering factor for early endosomal membranes ^18^, is localized to the somatodendritic region of hippocampal neurons ^19^ and therefore, cannot participate in axonal transport. Interestingly, the Rab5 effector APPL1 is an adaptor protein for Akt, a central kinase regulating cell survival ^20,21^ and plays a role in survival signaling from endosomes ^22^. APPL1 is localized to the dendritic spines and axons of hippocampal neurons ^23,24^. Its overexpression increases the amount of phospho-Akt (p-Akt) at synaptic sites, leading to formation of spines and an increase in synapse number ^24^. However, the data so far are restricted to immunocytochemistry and, there is no analysis of APPL1 endosome motility. Similarly, other Rab5 effectors have been poorly studied in neurons. Rabankyrin-5 is involved in macropinocytosis and is present in the growth cone of developing hippocampal axons but it is unclear whether it localizes to other neuronal endosomal compartments ^25^. Finally, the distribution of Rabenosyn-5, which plays a dual role in endocytosis and recycling ^13^ has not been addressed in neurons.

Here, we undertook a detailed comparison of the localization of endogenous Rab5 effectors and used live-cell imaging to quantitatively analyze the motility of axonal endosomes in pyramidal neurons, the most abundant type of neurons in the cerebral cortex and hippocampus.

## Results

### Rab5 displays complex motility in axons

Endosome motility and function is directed by Rab GTPases. In axons, it is unclear which Rab proteins play a role in signaling endosome motility although Rab5 labels NGF carrying endosomes in immunocytochemistry ^9^. Firstly, we aimed to characterize the motility of Rab5 endosomes in axons to determine if they display confined or long range movement, which is compatible with the transport of signaling endosomes, or a combination of both. We used fluorescent video microscopy to image primary mouse hippocampal neurons grown in a microfluidic chamber in which axons could grow up to 450 μm along the microgrooves separating two chambers ^26^. Neurons were plated on one side of the microfluidic chamber and agarose beads containing BDNF were deposited on the opposite side, in order to stimulate axonal growth. Axons emerging from cell bodies penetrate and grow along the microchannels until they reach the source of BDNF on the other side. The microfluidic chamber allows for the separation of axons and dendrites and facilitates imaging directional movement of endosomes in axons of hippocampal neurons. Using this setup, we imaged the axons of hippocampal neurons at day in vitro (DIV) 15 upon expression of mCherry-Rab5 via lentivirus transduction. We used a spinning disc microscope allowing for fast imaging of endosomes at frame-rate 2–4 frames per second (fps). According to our observations in other cell types ^15^, this is required to properly follow Rab5 endosomes moving at speeds higher than 2 μm/s that would otherwise escape detection. Endosomes were identified and tracked using the software platform Motiontracking^15,27,28^ and statistics were calculated both for object and track parameters including vesicle size, intensity, speed and total displacement as well as time spent on processive movement (consecutive steps in the same direction) and distance (see Methods).

We observed a variety of motility patterns in Rab5-positive endosomes ranging from stationary to highly processive (Fig.1A, B, C and movie 1). Displacement (distance of non-stop unidirectional movement) distribution of mCherry-Rab5 tracks showed a broad range with a mean displacement of 3.2±0.5 μm in retrograde and 1.9±0.4 μm in anterograde directions (Fig. 1D blue and red curves and Supp. Table 1). The mean represents the majority of the events, whereas the displacement spreads from 0.1 for slow-moving/non-processive all the way to 20 μm for the long-running endosomes. Rab5 endosomes moved processively in both anterograde and retrograde directions with similar displacement. Fig. 1B and Fig. 1C represent a kymograph from one axon at a time frame in which retrograde movement was more pronounced. We chose to represent retrograde movement in the kymograph because it correlates to the directionality of movement reported for signaling endosomes. Anterograde movement can be appreciated in Movie1, which represents the complete time series from which the kymograph was extracted. Although at low speed the distributions of anterograde and retrograde movements were almost identical, at high speed (> 1 μm/sec) the retrograde processive movement was much more pronounced (Fig. 1E, red anterograde and blue retrograde histograms). This results in a faster mean retrograde speed 0.82±0.12 μm/sec than the mean anterograde speed 0.53±0.04 μm/sec (Student’s-t test p_value_ = 0.037) (Supp. Table 1). To analyze this difference, we fitted each speed distribution (Fig. 1E red and blue solid lines) to two log-normal components. Both distributions share similar low speed components (Fig. 1E red and blue dotted line) but significantly different fast speed components (Fig. 1E red and blue dashed lines). Although both fast speed components have similar mean velocity (v_retrograde_ = 1.88±0.2 μm/sec and v_anterograde_ = 1.84±0.1 μm/sec, see details in legend of Fig. 1E), the retrograde constitute ~30% but the anterograde only ~2.5% of movement events. The values of fast speed components of processively moving Rab5 endosomes are in agreement with those of NGF transport in sensory and sympathetic axons, and sciatic nerve, which ranged between 0.56 and 5.6 μm/s with an average 1.75 μm/sec^7^.

**Figure 1:**
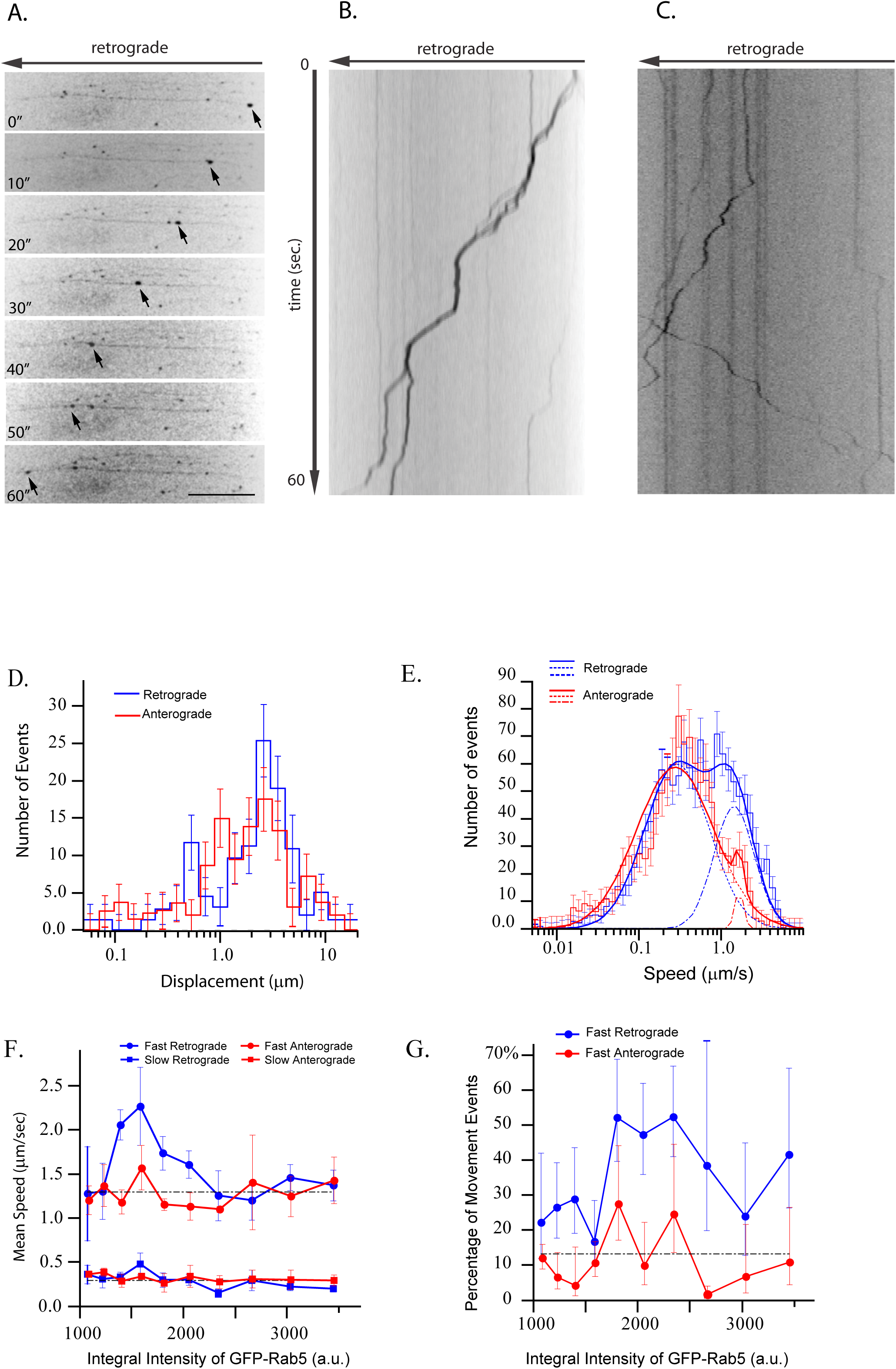
Rab5 endosomes display long-range movement. Primary hippocampal neurons were grown on a microfluidic chamber and transduced with mCherry-Rab5 lentivirus. Live cell imaging was performed at DIV 15. **(A)** Arrows point out the processive movement of a Rab5 labeled structure in a time series, bar 16.7 μm. **(B)** Kymograph shows the displacement of endosomes over time. Cell bodies are on the left side of the image. **(C)** Kymograph shows bi-directional movements of Rb5-positive endosomes in axon. **(D)** Distribution of displacements of mCherry-Rab5 labeled endosomes for the anterograde (red histogram) and retrograde (blue histogram) directions. **(E)** Speed distributions of mCherry-Rab5 labeled structures in the anterograde (red histogram) and retrograde (blue histogram) directions. Each distribution was decomposed into two log-normal components. Anterograde movement (red solid line) consists of two components, a slow component, μ=0.33 ± 0.014μm/sec, σ=0.96 ± 0.04, <v>=0.53 ± 0.03 μm/sec (dashed red lines) and a fast component, μ=1.8 ± 0.1μm/sec, σ=0.13 ± 0.05, <v>=1.84 ± 0.1 μm/sec (alternated dashed and dotted red lines). Similarly, retrograde movement (blue solid line) has a slow component, μ=0.33 ± 0.04μm/sec, σ=0.98 ± 0.08, <v>=0.53 ± 0.08 μm/sec (dashed blue lines) and a fast component μ=1.6 ± 0.2μm/sec, σ=0.57 ± 0.06, <v>=1.88 ± 0.19 μm/sec (alternated dashed and dotted blue lines). The fast anterograde and retrograde components constitute 2.5% and 30% of total movement events, respectively. The mean speeds for components were calculated by the formula 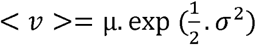. **(F)** Dependency of mean speed of the two log-normal components of endosome retrograde (blue) and anterograde (red) movement on (binned) integral intensity of Rab5, with slow components (square marked curves) and fast components (circle marked curves). Dashed black lines depict mean value of anterograde movement over all intensity bins. The fast retrograde speed is significantly different from the one of fast anterograde, Student’s-t p_value_=0.0014. **(F)** Dependency of fraction of movement events on (binned) integral intensity of GFP-Rab5 for fast anterograde (red) and retrograde (blue) motions. The t-test revealed significant difference between these dependencies (Student’s-t p_value_ = 0.004). Error bars indicate SEM. Statistical analysis was performed on 125 processive tracks from 4 movies from different experimental days.

To determine whether the motility correlates with the amount of Rab5 on endosomes, we grouped endosomes by GFP-Rab5 integral intensity and repeated the speed component analysis for each group (Fig. 1F, G). Slow speed components (both anterograde and retrograde) were independent of the amount of Rab5 on endosomes (Fig. 1F red and blue squares). Fast speed components showed differences with respect to the dependency on Rab5 on endosomes. Whereas the fast anterograde component was independent of the levels of Rab5 (Fig. 1F, red circles), the mean velocity of fast retrograde movement revealed a non-monotonic dependency on endosomal Rab5 (Fig. 1F, blue circles). A specific subpopulation of Rab5 endosomes displayed fast retrograde motility (Student’s-t p_value_=0.0014). The fast anterograde motility was not dependent on the amount of Rab5 (Fig. 1G, red line). In contrast, a large fraction (>50%) of endosomes with high GFP-Rab5 content displayed fast retrograde motility (Fig. 1G, blue line; Student’s-t p_value_=0.004). Altogether, our quantitative measurements suggest that the directionality of Rab5 endosomes in axons is biased toward the retrograde direction and fast retrograde movement depends on the levels of Rab5 on endosomes.

### Characterization of Rab5 effectors in hippocampal neurons

Rab GTPases orchestrate traffic through the recruitment of effector proteins to membranes. Many Rab5 effectors have been identified and serve different functions ^14^. The canonical Rab5 effector EEA1, important for tethering and fusion ^29^, is absent from axons ^19^. Thus, we wanted to systematically identify which other Rab5 effectors were present in axons in hippocampal neurons, in order to be eligible to act in signaling endosomes. APPL1 has been localized to axons and is an adaptor protein that binds to Akt ^23,30^. Another Rab5 effector, Rabankyrin-5, was found in growth cones of hippocampal neurons ^25^ suggesting that it may also be distributed in axons. Rabankyrin-5 is found on early endosomes and on macropinosomes, endocytic vesicles that take up fluid as well as protein aggregates, such as those generated during ALS and HD progression, and have been related to signaling endosomes via the Pincher protein ^31^. Additionally, Rabenosyn-5 has not been examined in hippocampal neurons, but there is evidence for a role of Rabenosyn-5 in recycling ^13^.

As we previously observed that overexpression or tagging proteins can yield localization and functional artifacts^15^, we used antibodies to detect the endogenous Rab5 effectors Rabankyrin-5, Rabenosyn-5, APPL1 and EEA1 in primary hippocampal neurons. Because all antibodies were rabbit polyclonals, the Rab5 effectors could be detected only individually without assessing their co-localization. In order to differentiate between axons and dendrites, we co-immunostained the Rab5 effectors with an axonal marker, phosphorylated neurofilament-1. For this set of experiments, neurons were grown following the Banker protocol ^32^. This culture set-up consists of the co-culture of astrocytes and neurons where astrocytes provide support for neuronal growth. This astrocyte feeder layer is compatible with low density seeding of neurons, which improves the identification of axons and other neuronal structures. As expected from previous findings ^19^, EEA1 labeled structures localized to the cell soma and dendrites and were absent from axons (Fig. 2B). Rabankyrin-5, Rabenosyn-5 and APPL1 localized to the cell soma and dendrites and were also present on axons (Fig. 2A, C and D). The EEA1 puncta in the cell body were larger than the APPL1 and Rabenosyn-5 puncta, and similar in size to the structures harboring Rabankyrin-5 (that had a high cytosolic pool and, therefore, whose puncta in axons were more difficult to resolve). In addition, a fraction of APPL1 was found in the nucleus, as previously shown for other cell types ^33^. The subcellular distribution suggests a role for Rabenosyn-5, Rabankyrin-5 and APPL1 in membrane traffic/signaling events in axons. Among the Rab5 effectors, APPL1 was previously shown to interact with signaling receptors and with the three isoforms of Akt/PKB in various cell system^23,30,34–36^. Given that APPL1 is localized to axons and can be tagged without compromising its localization and function of the endosomal system ^15^, we decided to focus on this protein as a candidate Rab5 effector on a signaling endosome.

**Figure 2:**
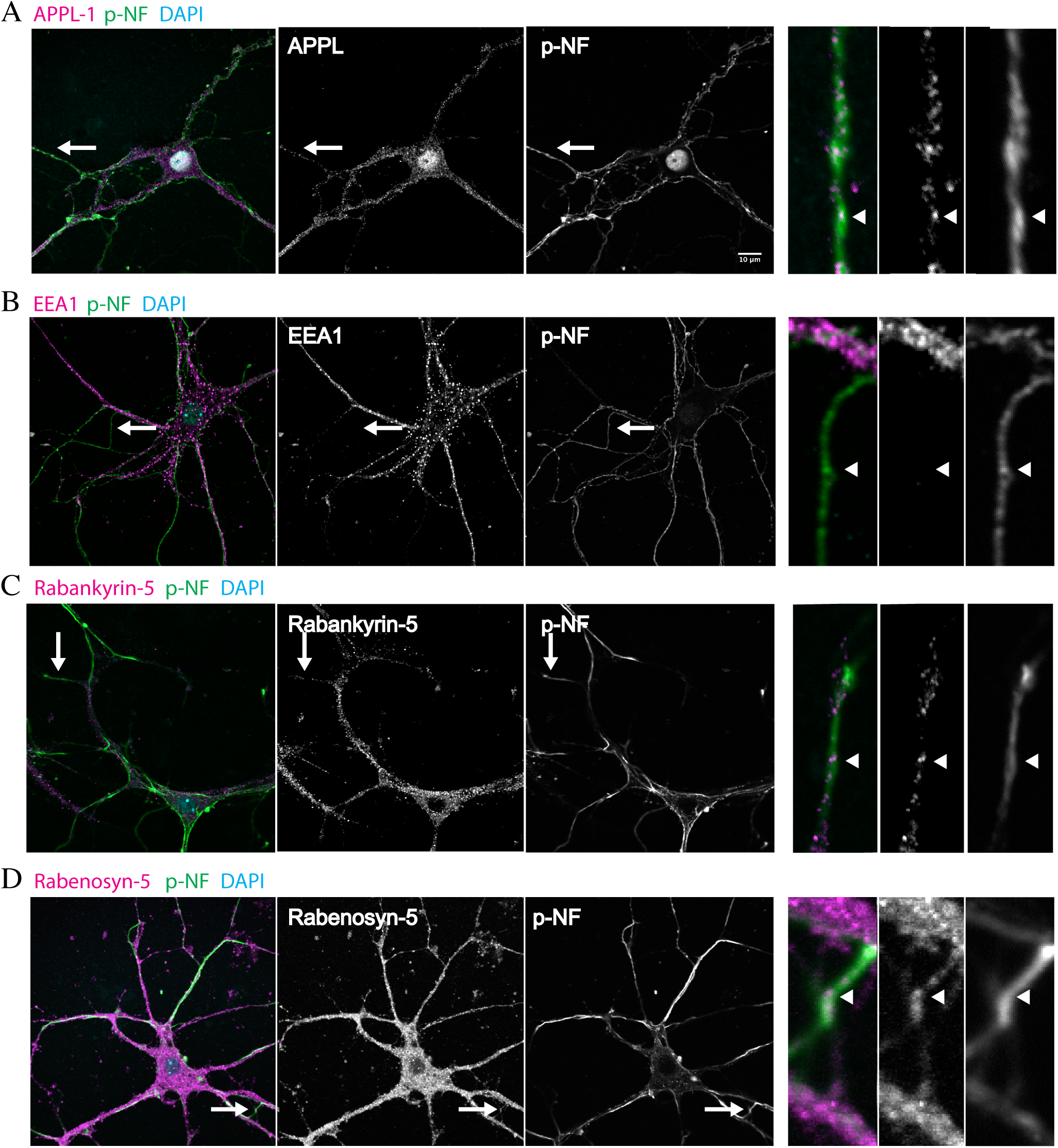
Rab5 effectors localize to axons and dendrites. Primary hippocampal neurons were grown at low density supported by an astrocyte feeder layer. At DIV 14, neurons were fixed and immunostained for phosphorylated neurofilament-1 (pNF) and APPL1 **(A)**, EEA1 **(B)**, Rabankyrin-5 **(C)** or Rabenosyn-5 **(D)**. The inset highlighted by the arrow shows a respective axon whereupon there is co-localization of APPL1, Rabankyrin-5 and Rabenosyn-5 with pNF but not with EEA1. Bars 10 μm.

### APPL1 structures move bi-directionally with significant retrograde bias

We first validated the localization of APPL1 to Rab5 endosomes by co-expressing GFP-Rab5 and mCherry-APPL1 (validated as in ^15^), and performing live cell imaging in hippocampal neurons (Movie 2). APPL1 motility was assessed using conditions similar to the aforementioned Rab5 movies (Fig. 3A, B and Movie 3, Supp. Fig. 1 and Supp. Movie 1). In axons, mCherry-APPL1 endosomes displayed long-range movements with a mean track displacement of 1.70 ± 0.04 μm in the retrograde and 1.46 ± 0.04 μm (mean±SEM) in the anterograde direction (Fig. 3C and Supp. Table 1; Student’s-t p_value_ = 5·510^−5^). Average speeds of APPL1 endosomes in both retrograde and anterograde directions were equivalent (0.35±0.04 μm/sec and 0.33±0.04 μm/sec, respectively). Analysis of log-normal components (Fig. 3D blue and red dashed lines) revealed that retrograde and anterograde speed distribution share similar slow components (0.23±0.01 μm/sec and 0.24±0.01 μm/sec). Surprisingly, the fast speed component had significantly higher mean value in the anterograde (1.84±0.1 μm/sec) than in the retrograde (1.2±0.1 μm/sec) direction (see Fig. 3D, dashed red and blue curves and tracks marked by blue and red arrows on Supp. Fig. 1A, B). However, the fraction of fast retrograde movement was almost 3 fold larger (17±1.5%) than the anterograde one (6±1.5%), resulting in a net retrograde flux of APPL1-positive endosomes. This is consistent with the behavior of a sub-population of Rab5 endosomes (Fig. 1E).

**Figure. 3:**
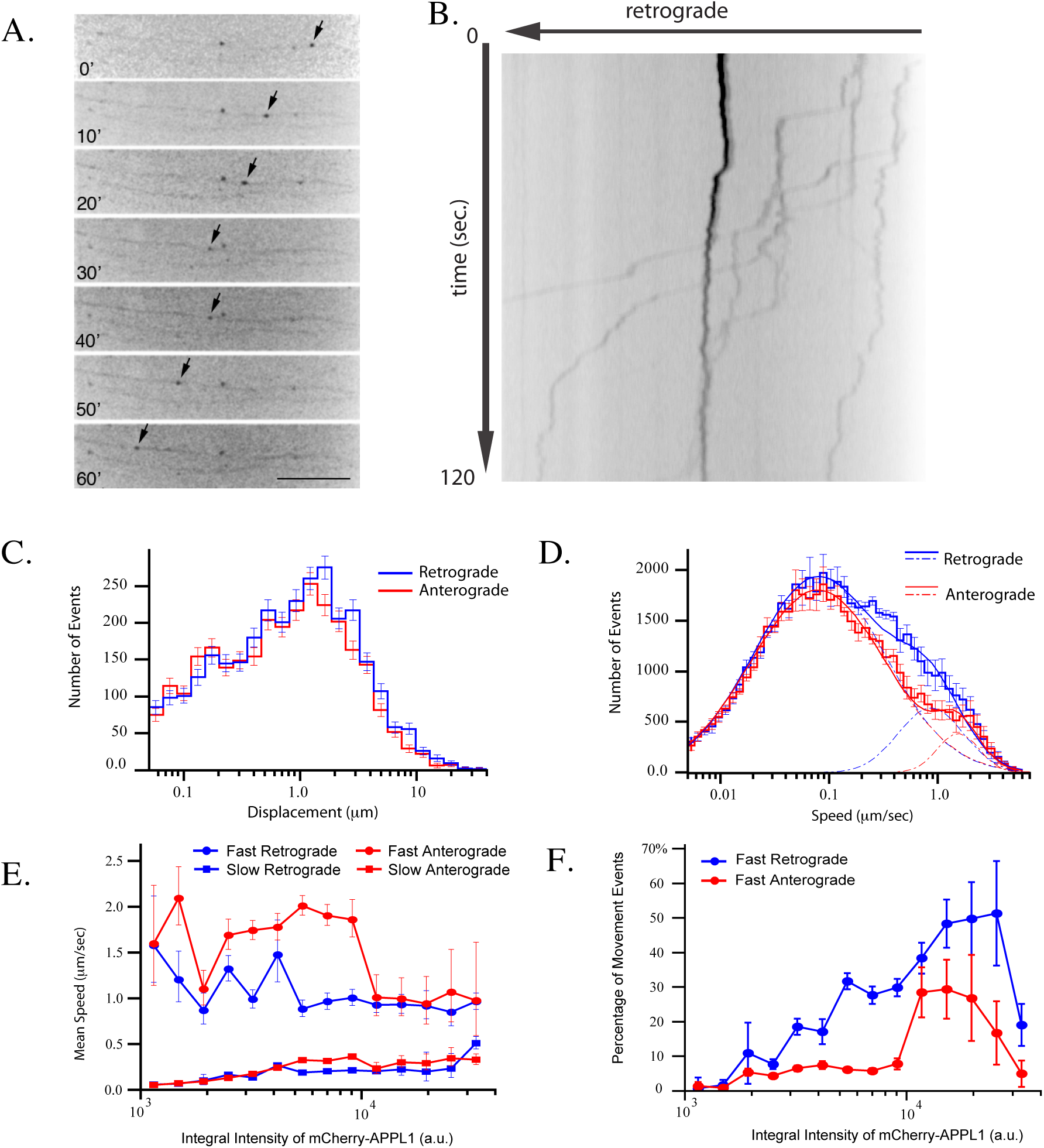
APPL1 localization to endosomes is correlated to speed. Primary hippocampal neurons were grown and imaged as in Fig. 1. **(A)** Arrows point out the processive movement of one APPL1 labeled structure in a time series, bar 16.7 μm. Cell bodies are on the left side of the image. **(B)** Kymograph shows the displacement of endosomes over time. Statistical analysis was performed on 1603 tracks from 6 movies within the chamber from different experimental days. **(C)** Displacement distribution of mCherry-APPL1 labeled structures moving in anterograde (red histogram) and retrograde (blue histogram) direction. **(D)** Speed distributions of mCherry-APPL1 labeled structures in the anterograde (red histogram) and retrograde (blue histogram) directions. Anterograde movement (red solid line) consists of two components, a slow component, μ=0.085 ± 0.002μm/sec, σ=1.4. ± 0.03, <v>=0.24 ± 0.01μm/sec (dashed red lines) and fast component, μ=1.67 ± 0.1μm/sec, σ=0.43 ± 0.04, <v>=1.84 ± 0.1μm/sec (alternated dashed and dotted red lines). Retrograde movement (blue solid line) has two components, which parameters are: slow component, μ=0.087 ± 0.002μm/sec, σ=1.38 ± 0.02 (dashed blue lines) and fast component μ=0.94. ± 0.05 μm/sec, σ=0.69 ± 0.03, <v>=1.2 ± 0.1 μm/sec (alternated dashed and dotted blue lines). The fast anterograde and retrograde components constitute 6 ± 1.5% and 17 ± 1.5% of total movement events, respectively. **(E)** Dependency of mean speed of the two log-normal components of endosome retrograde (blue) and anterograde (red) movement on (binned) integral intensity of mCherry-APPL1, with slow components (square marked curves) and fast components (circle marked curves). Dashed black lines depict mean value of retrograde movement over all intensity bins. The fast retrograde speed is significantly different from the one of fast anterograde, Student’s-t p_value_=0.0004. **(F)**. Dependency of fraction of movement events on (binned) integral intensity of mCherry-APPL1 for fast anterograde (red) and retrograde (blue) motions. The t-test revealed significant difference between these dependencies (Student’s-t p_value_ = 0.0004). Arrowheads: retrograde movement; bars 10 μm. Error bars indicate SEM.

Next, we analyzed the dependency of speed on the amounts of APPL1 on endosomes. Interestingly, it was drastically different from that of Rab5. Anterograde and retrograde slow components had similar speed and positive correlation with the levels of APPL1 (Fig.3E, red and blue rectangles). The mean speed of the fast retrograde component was independent of the levels of Rab5, whereas the anterograde component displayed a negative correlation with the amounts of APPL1 on endosomes (Fig. 3E, red and blue circles, Student_t p_value_=0.0004).

We also analyzed the fraction of fast retrograde motion events associated with the amounts of APPL1 and observed that it increased from 5 to 60% with the amounts of APPL1 on endosomes (Fig. 3F blue circles). In contrast, the fraction of fast anterograde motion events was ~ 5% and was independent of the amount of APPL1 up to some threshold (~10^4^ fluorescence a.u.). Above such threshold the fraction of fast movements increased up to 30% (Fig.3F, red and blue curves). Altogether, these results indicate that both Rab5 and APPL1 endosomes move more frequently and processively in the retrograde than in the anterograde direction. However, the levels of APPL1 correlate more with fast retrograde movement than the levels of Rab5, arguing that they mark a population of fast retrogradely moving Rab5 endosomes.

### APPL1 endosomes transport signaling molecules

The preferential retrograde motility of APPL1 endosomes prompted us to ask whether they transport signaling cargo. To address this question, we co-expressed GFP-TrkB with mCherry-APPL1 and performed dual-color simultaneous imaging. The motility of APPL1- and TrkB-positive endosomes is presented on Fig. 4A and Movie 4. A significant fraction (33±5%) of GFP-TrkB carrying endosomes were mCherry-APPL1 positive. However, this is in all likelihood an under-estimate, as the mCherry-APPL1 signal was much dimmer than that of GFP-TrkB. APPL1 endosomes transported TrkB bi-directionally (blue and white arrows on Fig. 4A). We found that mCherry-APPL1 unevenly distributed between retrograde and anterograde moving TrkB-positive endosomes. The fraction of retrograde moving TrkB positive endosomes with detectable APPL1 (29±6%) was significantly larger (Studet_t p_value_ = 0.02) than the fraction of anterograde moving APPL1- and TrkB-positive endosomes (11±2.5%). Although some anterograde movements were fast and processive, the net flux of GFP-TrkB-endosomes was retrograde. Interestingly, the amount of APPL1 on TrkB-positive signaling endosomes was not constant and the intensity variations correlated with endosome speed (white arrowheads on Fig 4A). The speed distributions of anterogradely (red curves) and retrogradely (blue curves) moving TrkB-positive endosomes were similar to those of APPL1 endosomes (Fig. 4B, compare with fig. 3D). Decomposition of the curves shows that the slow moving endosomes had similar velocities for anterograde (0.31±0.025 μm/sec, dashed red lines) and retrograde (0.36±0.04 μm/sec, dashed blue lines) movements. High speed movement components on the anterograde direction were faster (2.5±0.45 μm/sec, dashed-dotted red lines) than retrograde movements (1.47±0.3 μm/sec, dashed-dotted blue lines). However, the total retrograde flux (product of speed by number of movement evens) was ~25% larger than the anterograde flux. The speeds of double positive APPL1 and TrkB endosomes were slower than those endosomes with no or faint mCherry-APPL1 signal (mean retrograde speed 0.24±0.04 μm/sec and anterograde speed 0.21±0.06 μm/sec), presenting only a slow speed component (Fig. 4C). The speed of slow component does not dependent on direction. However, there are much more double positive endosomes which move retrogradly than anterogradely (331 anterograde events vs. 812 retrograde events). We addressed the correlation between the amount of APPL1 on TrkB endosomes and endosomal speed and observed that the velocity slightly decreases with increase APPL1 on endosomes (Fig. 4D). Correlation of number of movements and APPL1 integral intensity, shows that there is no dependency of the number of retrograde events on APPL1 amounts on endosomes but the number of anterograde events significantly decrease with increase amount of APPL1 (Supp. Fig. 2). From these data, we conclude that APPL1 endosomes carry a fraction of axonal transported TrkB which present lower speed movements, move more frequently in the retrograde direction.

**Figure 4:**
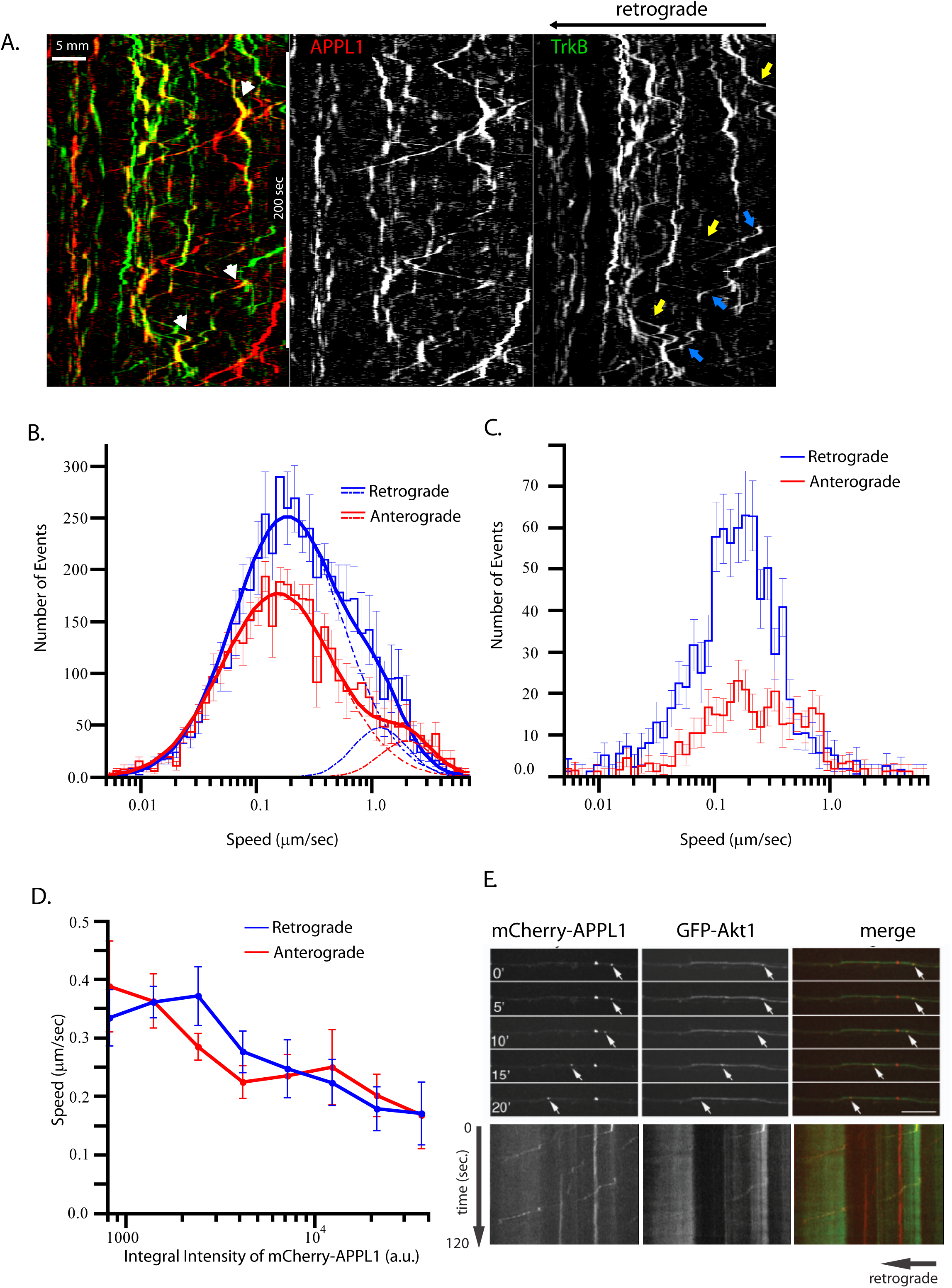
APPL1 endosomes transport signaling cargo. Primary hippocampal neurons grown on a microfluidic chamber were transduced with mCherry-APPL1 and GFP-TrkB lentivirus. All live cell imaging experimens were performed at DIV 15. **(A)** Kymographs representing movie 4. Cell bodies are localized towards the left side of the image. Yellow and blue arrows on TrkB panel point to anterograde and retrograde directed tracks respectively. White arrow heads on merge image points on APPL1 fluctuation on APPL1- and TrkB-double-positive tracks. **(B)** Speed distributions of GFP-TrkB labeled structures in the anterograde (red histogram) and retrograde (blue histogram) directions. Anterograde movement (red solid line) consists of two components, a slow component, μ=0.17 ± 0.01 μm/sec, σ=1.1 ± 0.04, <v>=0.31 ± 0.02 μm/sec (dashed red lines) and a fast component, μ=2.16 ± 0.3 μm/sec, σ=0.53 ± 0.12, <v>=2.5 ± 0.45 μm/sec (alternated dashed and dotted red lines). Retrograde movement (blue solid line) has a slow component, μ=0.2 ± 0.02 μm/sec, σ=1.08 ± 0.06, <v>=0.36 ± 0.04 μm/sec (dashed blue lines) and a fast component μ=1.27 ± 0.24 μm/sec, σ=0.53 ± 0.14, <v>=1.5 ± 0.3 μm/sec (alternated dashed and dotted blue lines). **(C)** Speed distributions of GFP-TrkB and APPL1 double labeled structures in the anterograde (red histogram) and retrograde (blue histogram) directions. **(D)** Dependency of slow component velocity of double-positive mCherry-APPL1 and GFP-TrkB endosomes on integral intensity of mCherry-APPL1 per endosome for retrograde (blue) and anterograde (red) directions. **(E)** Primary hippocampal neurons grown on a microfluidic chamber were transduced with mCherry-APPL1 and GFP-Akt lentivirus. Time series and corresponding kymograph are shown. Bars 10 μm.

Since APPL1 was reported to interact with Akt we investigated whether APPL1 endosomes are able to recruit Akt1 and transport it retrogradely. The largest pool of Akt1 is cytosolic, making it difficult to monitor the endosomal pool. To overcome this problem, we performed fast dual-color simultaneous imaging of neurons expressing mCherry-APPL1 and GFP-Akt1 and applied the de-noising algorithm *BFBD* ^37^ to suppress noise and Frog-Eye Filter (http://motiontracking.mpi-cbg.de) to reveal moving endosomes in high cytoplasmic background (compare Supp. Fig 1A-B and Supp. Movie 1 and Supp. Movie 2). Under our experimental conditions, we were able to detect Akt1- as well as APPL1-positive endosomes moving along axons (Fig.4E). By comparing APPL1 anterograde and retrograde movement speed with and without GFP-Akt expression, we determined that when GFP-Akt is co-expressed with APPL1, anterograde speed remains unchanged whereas retrograde speed increases. In addition, in the presence of Akt, long-range movement occurs with faster kinetics (supp. Table 1), suggesting that Akt regulates endosome motility. Since the de-noise approach was not sufficient to determine the amounts of Akt on APPL1 endosomes we decided to perform FRET/FLIM experiment to further investigate the Akt/APPL1 interaction.

### Akt1 interaction with APPL1 is dependent on endocytosis

Our data suggest that Akt1 is transported retrogradely in axons in Rab5 and APPL1-positive endosomes. To determine whether APPL1 and Akt1 interact on axons, we used a FRET (Fluorescence energy transfer)/FLIM (fluorescence lifetime imaging) approach to more precisely analyze protein-protein interactions compared to conventional light microscopy (resolution of 8 nm and 200 nm, respectively). In this assay, two distinct fluorescent proteins transfer energy from one to another, one acting as a donor and another as an acceptor. When energy transfer occurs in the presence of an acceptor, the lifetime of the donor is shortened. Lifetime measurements can be converted to FRET efficiency values using the equation *E* = 1 – (τ_DA_/τ_D_). In our experiments, we used GFP-Akt1 as a donor and mCherry-APPL1 as the acceptor as FRET pair ^38^. Lifetime of the donor was measured in the absence (τ_D_) and in the presence of acceptor (τ_DA_).

To measure the interaction between APPL1 and Akt1 using FRET, we analyzed live neurons expressing GFP-Akt1 alone and the pair GFP-Akt1/mCherry-APPL1. Interestingly, we observed that the lifetime of GFP-Akt1 was significantly decreased in the growth cone but not in the soma of neurons co-expressing mCherry-APPL1 in comparison with neurons expressing GFP-Akt1 alone (Fig. 5A) (exp1 n=10; exp2 n=10). This shows that Akt1 interacts more with APPL1 in axons than in the soma as reflected by the higher FRET efficiency values in the growth cone than in soma (Fig. 5B). To confirm the specificity of the FRET signal observed, we used an APPL1 mutant devoid of the Akt1 interaction site, i.e. a truncated version lacking the PTB domain ^30^. As expected, the lifetime of GFP-Akt1 did not change in the presence of the acceptor mCherry-APPL1-delta-PTB resulting in FRET efficiencies that corresponded to zero in the cell body or values lower than 1% in the growth cone (Fig. 5B). This indicates that the decrease in the lifetime of GFP-Akt1 is specific to the APPL1-Akt1 interaction.

**Figure 5:**
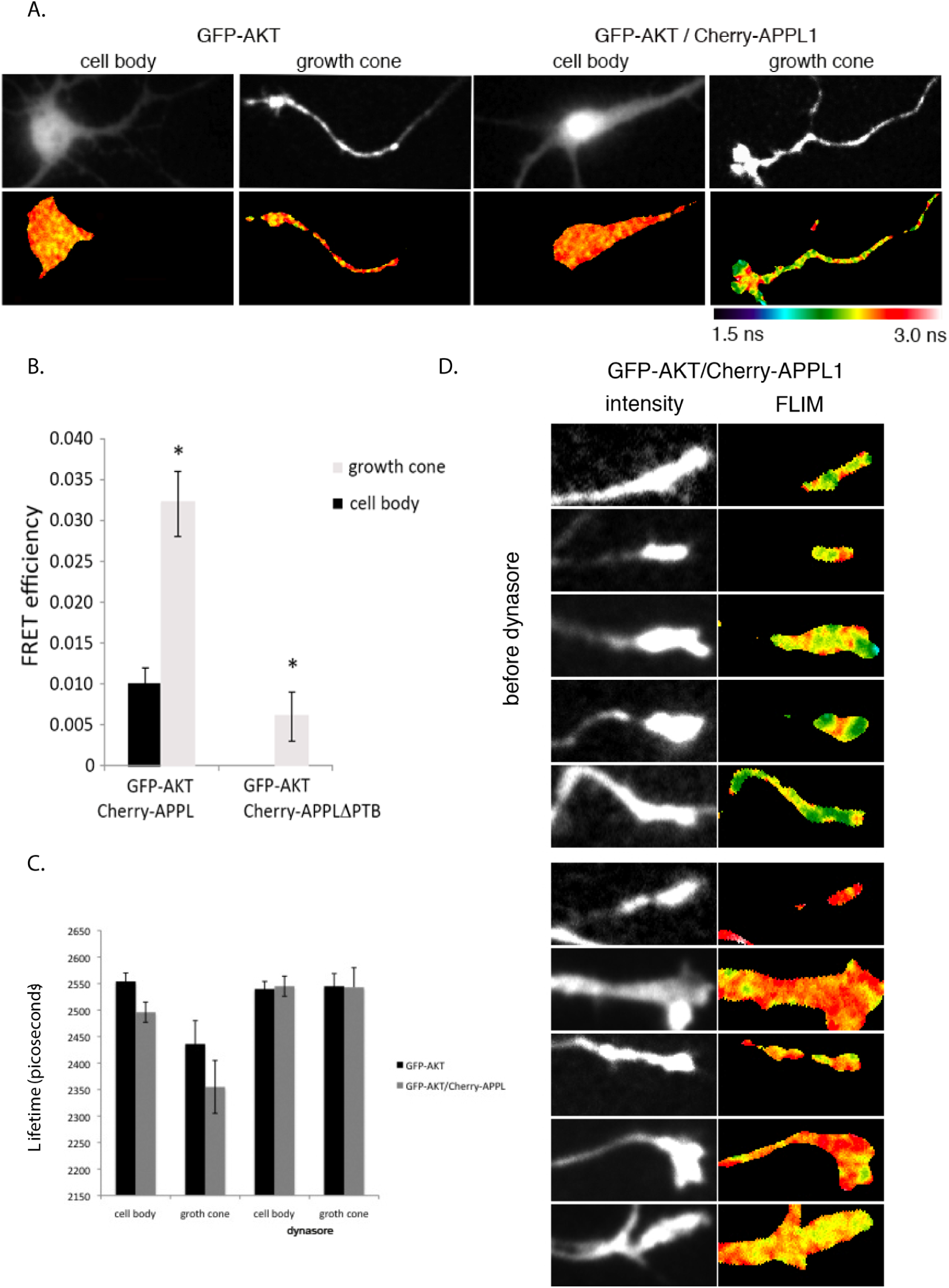
APPL1 and Akt1 interact at the neuronal growth cone. Hippocampal neurons were plated at low density and transduced with lentivirus to express the donor GFP-Akt1 alone or in the presence of the acceptor protein fused to mCherry, mCherry-APPL1 or mCherry-APPLΔPTB a truncated form of APPL1 lacking Akt binding domain. At DIV 1, neurons were transferred to an astrocyte feeder layer. Live cell FLIM measurements were performed at DIV 7. **(A)** Neurons expressing GFP-Akt1 alone display higher GFP lifetimes (yellow-red range) when compared to neurons expressing GFP-Akt1 and mCherry-APPL1 (green-blue range). The decrease in lifetime of GFP-Akt is evident in the axons. **(B)** FRET efficiency at the cell body and growth cones when GFP-Akt1 was co-expressed with mCherry-APPL or with mCherry-APPLΔPTB a truncated form of APPL1 lacking Akt binding domain. Error bars represent SEM from 3 independent experiments totalizing 60 cell measurements (*p<0.001). **(C)** Dynasore treatment abolishes the decrease in lifetime of GFP-Akt1 in the presence of mCherry-APPL1. **(D)** Lifetime measurements of GFP-Akt1 alone or in the presence of mCherry-APPL1 before and after dynasore treatment. Error bars represent SEM from 3 independent experiments totalizing 90 cell measurements.

Signaling endosomes receive cargo and many growth factor receptors internalized via Clathrin- and Dynamin-mediated endocytosis. Since the small molecule dynasore ^39^ blocks Dynamin-dependent endocytosis in neurons ^40^, we used it to test the requirement of endocytosis for the APPL1-Akt1 interaction. We performed the FRET/FLIM assay in neurons before and after dynasore treatment for 5 min. We observed no APPL1-Akt1 interaction in growth cones after dynasore treatment (Fig. 5C). Lifetime measurements of cell bodies and growth cones of neurons showed that the lifetime of GFP-Akt1 was lower in the presence of mCherry-APPL1. After dynasore treatment, mCherry-APPL1 was no longer able to reduce GFP-Akt1 lifetime, indicating that under these conditions the two proteins do not interact (Fig. 5D). These data suggest that endocytosis is required for the interaction of Akt1 and APPL1, thus providing further evidence that axonal retrograde APPL1 vesicles may be signaling endosomes.

## Discussion

In this study, we report a detailed quantitative characterization of Rab5 early endosome motility in axons of primary hippocampal neurons. This analysis has revealed that Rab5 endosomes follow a complex pattern of motility that can be ascribed to different populations of endosomes differing in directionality, speed and processivity. These populations correspond to endosomes with different content of Rab5, the Rab5 effector and signaling adaptor APPL1, and the signaling kinase Akt1. Among the Rab5 endosomes, the fraction that moves retrogradely with higher frequency than in the anterograde direction, speed and processivity corresponds to a sub-population of organelles harboring APPL1. Such endosomes may participate in neurotrophin retrograde movement and signaling. Long range movement of Rab5 endosomes was also observed in motor neurons at speeds compatible to that reported for neurotrophin transport (data not shown). Consistent with this idea, APPL1 was co-transported with TrkB and Akt on endosomes, suggesting that it could bridge receptors to signaling molecules. Akt co-expression stimulated retrograde movement velocity and processivity. Finally, we found that endocytosis is required for the interaction of Akt1 and APPL1, thus providing further evidence that axonal retrograde APPL1 vesicles may be signaling endosomes.

The molecular identity of signaling endosomes in neurons has been controversial. In motor neurons, Rab7 exhibited retrograde movement whereas Rab5 was reported to remain stationary ^12^. On the other hand, Rab5 was found in signaling endosomes collected from sciatic nerve ^7^ and Rab5 endosomes transport NGF in dorsal root ganglia ^9^. NGF transport via Rab5-positive endosomes has been previously visualized indirectly using quantum-dots labeled NGF ^9^. However, due to their size quantum dots can interfere with the normal receptor trafficking, by triggering the internalization of receptors and routing them toward endosomal degradation ^41^. Our observations shed new light on this problem. The velocity of Rab5 endosome movement visualized in non-neuronal cells ^15^ and in the axon shown here is such that requires imaging at high frame rate (≥ 2 fps) and overcoming technical challenges posed by high background of soluble markers proteins. The discrepancy between the published studies showing only oscillatory Rab5 movement^12^ and ours could be explained by neuronal subtype difference, cargo specificity, overexpression conditions or differences between the imaging set up used. For example, acquisition at lower frame rates (0.2 fps)^12,64^ and lower signal-to-noise ratio reported in previous studies certainly compromise the ability to track very fast and dim objects (Fig. 1F, Fig. 4E, Supp. Fig. 1), in addition, low throughput and manual analysis^64^ could prevent detecting the retrograde bias of bi-directional motility of Rab5-positive endosomes (Fig. 1E-G). Nevertheless, it is possible that signaling molecules can be transported non-exclusively, i.e. by different endosomal types. In hippocampal neurons, multivesicular bodies (MVB) transport endocytic cargo retrogradely in the axon ^10^. This is in line with a recent study showing that MVB transport NGF-TrkA between the nerve terminals and the cell body in sensory neurons ^42^. In sympathetic neurons Rab7 and Rab11 endosomes carry TrkA^43^. We still cannot exclude the possibility that the Rab5 endosomes visualized in axons could be Rab5-to-Rab7 conversion intermediates ^27^. However, two arguments are against this interpretation. First, the reported time for Rab5-to-Rab7 conversion of endosomes is much shorter (0.5÷3 min) ^56, 63^ than the retrograde moving time of Rab5-positive endosomes in axons (> 10 min). Second, APPL1 is a Rab5 but not a Rab7 effector^15,16,33^ and mostly absent from the EEA1-positive endosomes capable for Rab5 to Rab7 conversion^15^. Unfortunately, we could not unambiguously exclude the presence of Rab7 by transiently co-expressing and imaging fluorescently tagged Rab5 and Rab7. Addressing this question requires validated genome-edited cells.

The different populations of motile Rab5 endosomes could also harbor Rab5 effectors other than APPL1. Whereas EEA1 is exclusively localized to the somato-dendritic compartment ^19^ and its localization is defined by the Rab5 effector motor protein KIF16B^41^ which regulates plus end motility of Rab5 endosomes^42^, Rabenosyn-5 and Rabankyrin-5 are also present in axons (Fig. 2). This suggests a differential distribution and function of Rab5 effectors on early endosomes within neurons. The underlying molecular mechanisms are unclear. Rabankyrin-5, Rabenosyn-5 and EEA1 all contain a FYVE domain which, by binding to phosphatidylinositol 3-phosphate (PI(3)P) and, together with binding to Rab5, supports recruitment to PI(3)P-positive early endosomes ^14,29,44^. These endosomes are present on axons where they recruit Ankyrin B in a PI(3)P-dependent manner^45^. Yet, the presence of FYVE domains and Rab5-binding sites are not sufficient to recruit EEA1 to these endosomes. Evidently, additional molecular interactions must play a role in the selection of Rab5 effectors present in the axon. Rabankyrin-5 participates in macropinocytosis, a process that has been related to NGF uptake and retrograde transport mediated by the pinocytic chaperone Pincher ^31^. Additionally, it has been linked to retrograde transport from endosomes to the Golgi complex via interaction with the retromer protein EHD1 ^46^. Rabenosyn-5 links early endosomes and the recycling route via its interaction with Rab5 and Rab4 ^13^. Given that endocytic/recycling regulates axon elongation, it is possible that Rabenosyn-5 plays a role in this process ^47^. In contrast to the aforementioned Rab5 effectors, APPL1 does not have a FYVE finger and is recruited to membranes via its BAR-domain and the interaction with Rab5.

Rab5-APPL1 endosomes display characteristics of neuronal signaling endosomes. APPL1 has molecular features of a signaling adaptor ^22,24,48–50^. It is transported together with TrkB and Akt on endosomes in a retrograde direction, suggesting that it could bridge receptors to signaling molecules. Although there have been demonstrations of ligand receptor complexes, phosphorylated substrates and accessory proteins moving retrogradely, to our knowledge, this is the first data showing the live movement of a kinase, along with the molecule that links it to endosomes in axons. Akt regulates axonal growth, regeneration and protects axons from degeneration ^51–53^. Endosomal transport may be crucial for these functions. Interestingly, overexpression of Akt affects differently APPL1 anterograde and retrograde movement speed (anterograde speed remains unchanged whereas retrograde speed increases), and increases long-range movement with faster kinetics (supp. Table 1). These results argue that Akt and perhaps other signaling molecules modulate the speed and processivity of the endosomal compartments harboring them. Interaction with Akt1 is dependent on the APPL1 PTB domain and is sensitive to pharmacological inhibition of endocytosis. This establishes a link between APPL1/Akt interaction and endocytosis. We did not expect changes on global Akt phosphorylation levels on the basis of its interaction with APPL1, since only a small fraction of Akt is concentrated on endosomes in comparison with the magnitude of the soluble pool. However, the endosomal fraction must be very important, since inhibition of endocytosis with dynasore impairs Akt phosphorylation ^54^. Therefore, it is conceivable that Akt phosphorylation relates to its interaction with APPL1 on endosomes. It was previously reported that TrkB colocalizes to APPL1 positive endosomes ^55^. Here, we showed that these endosomes undergo bi-directional movements with a strong bias in the retrograde direction. The fluctuations of the levels of Rab5 on endosomes correlate with cargo processing ^27,56^. We detected similar behavior for APPL1 on TrkB carrying endosomes with respect to retrograde speed modulation (Fig. 4A-C). It is known that the APPL1/Akt signaling complex regulates synapse formation ^24^ and BDNF. Neutrotrophins such as BDNF modulate synapse transmission, plasticity and growth ^57^. We propose that such functions may be mediated by the APPL1/Akt complex on endosomes that transit retrogradely from axons to the cell body to deliver signals for synapse regulation. Further molecular characterization of the APPL1/Akt endosomes, such as identification of signaling receptors, their ligands, the presence TrkB-BDNF complex as well as downstream signaling components are still required to gain a better understanding of the role of Rab5/APPL1/Akt endosomes in neuronal signaling.

The existence of APPL1-Akt signaling endosomes raises the question of whether these endosomes and downstream signaling are affected in disease. Alzheimer’s disease (AD) and other tauopathies are interesting candidates to investigate, given that they correlate with defects in the endolysosomal system ^6^. It has been hypothesized that AD is caused by hyperactivation of tau by GSK-3β, a kinase downstream of Akt ^58^. Akt phosphorylates and inactivates GSK-3β and our previous data suggest that APPL1 influences Akt activity towards GSK-3β, promoting its inactivation at membranes ^22^. In this sense, an increase in APPL1-Akt endosomes could have a neuroprotective effect in disease. Further exploring the role of Akt/GSK-3β activation at endosomes could contribute to the development of drugs aimed at specifically modulating GSK-3β, providing new opportunities to treatment of brain diseases.

## Material and methods

### Antibodies

Primary antibodies utilized in this study were: rabbit anti-human APPL1 (1:1000 in IF)^33^, rabbit anti-EEA1 (1:1000 in IF) ^33^, rabbit anti-Rabenosyn-5 (1:1000) ^13^, rabbit anti-Rabankyrin-5 (1:1000) ^44^, rabbit anti-Akt (Cat.# 9272, Cell Signaling, Beverly, MA, USA, 1:100 in IF), mouse anti-MAP2 (Cat.# MAB3418, Chemicon Europe, 1:1000 in IF), mouse anti-phosphorylated neurofilament antibody (Cat.# SMI-31, HISS Diagnostics, Freiburg, DE, 1:3000 in IF). Antibodies produced in house have been affinity-purified and extensively validated in other cells ^13,15,33,44^ Their specificity has been further demonstrated by the disappearance of the signal upon knock-down by RNAi. Their staining has been optimized in cultured neurons. Secondary antibodies used were all AlexaFluor conjugates from Invitrogen (1:1000 in IF).

### Primary culture of neurons

Hippocampal neurons were isolated from mouse embryos E14–17. All animal experiments were performed in accordance with German animal welfare legislation, guidelines and regulations and were approved by the Landesdirektion Sachsen (license no. DD24–9168.24–9/2009–1). Mouse hippocampi from embryos of either sex were dissected in phosphate buffered saline - PBS (25 mM Na-phosphate buffer, pH 7.4, 110 mM NaCl, 1 mM EDTA) and dissociated in digestion solution (100 mg/ml DNAse I and 200 Units Papain in PBS), for 20 min. Hippocampi were washed twice with plating media (DMEM containing 10% FCS, 2 mM glutamine, 50 mg/ml penicillin / streptomycin, Invitrogen, Karlsruhe, DE). Then, tissue was triturated in plating media and counted for plating. Neurons were plated in glass coverslips coated with 1 mg/ml poly-L-lysine (Cat. n^o^ P1524, Sigma-Aldrich Chemie GmbH) at high density (200,000 cells/ml) or at low density (25,000 cells/ml) in the presence of an astocyte feeder layer. The day after plating media was changed to Neurobasal medium with supplements (50 IU/50 mg/ml penicillin/streptomycin, 2 mM glutamine, 0.23 mM, 2% B27, Invitrogen).

Mouse astrocytes to be used as a supporting culture for the hippocampal neurons were obtained from mouse cortex dissected from mice of age P0-P3 of either sex ^32^.

After dissection, tissue was digested in 12 ml PBS containing 2 mg/ml trypsin, 0.1 mg/ml of DNAse for 5 min. After that the tissue was triturated by pipetting and further incubated in the digestion solution for 10 min. The tissue was triturated by pipetting again and passed through a 70 mm cell strainer (Cat. n^o^ 9663160, Fisher Scientific GmbH, Nidderau, DE) into tube with 15 ml of Plating Medium. Cells were spun down, resuspended, counted and finally plated at a density of 500,000 cells/ml using plating medium. The day before hippocampal neurons were inverted in the astrocyte feeder layers the medium of the astrocytes was changed to Neurobasal medium with supplements.

Neurons for plating in the microfluidic chamber were subjected to an OptiPrep density 1.32 (Cat. n^o^ D1556, Sigma) gradient to remove cell debris ^59^. Microfluidic chambers (Xona Microfluidics LLC, Aliso Viejo, CA, USA) ^26^ were mounted in a glass coverslip coated with 1 mg/ml Poly-L-Lysine. Neurons were plated on two wells on one side of the chamber and on the opposite side of the chamber recombinant human BDNF (Cat. n^o^ B-250, Alomone Labs, Jerusalem, Israel) diluted in agarose was added.

### Virus production and transduction

mCherry-Rab5, GFP-Rab5, GFP-TrkB, GFP-C3-G6-Akt1, mCherry-C3-APPL1 and mCherry-C3-ΔPPL-APTB were cloned into the lentivector pRRL.SF.pSIN.ppt kindly provided by Prof. Sebastian Brenner (Klinik für Kinder- und Jugendmedizin, Haus 21 Universitätsklinikum Carl Gustav Carus, Dresden). Cloning was done by restriction digestion of the lentivector with appropriate restriction enzymes and ligation with a restriction fragment obtained from another vector or from a PCR product. Lentivirus particles were produced in HEK 293T cells by co transfection of the lentivector and the accessory plasmids pPAX and pVSV-G using Polyethylenimine (PEI)(Cat. n^o^ 408727, Sigma-Aldrich Chemie GmbH, Taufkirchen, DE). HEK 293T cells were maintained in IMDM containing 10% (v/v) fetal calf serum (FCS), 2 mM L-glutamine, 100 U/ml penicillin and 100 μg/ml streptomycin (Invitrogen) under 5% CO_2_ in a humidified atmosphere at 37°C. The transfection media consisted in 10 mg of lentivector, 6.5 mg of pSPAX and 1 mg of pVSV-G diluted in 525 ml of IMDM to which 37.5 mg PEI in 525 ml of IMDM was added and incubated for 30 min. This mixture was resuspended to 5 ml with IMDM containing 10% (vol/vol) foetal calf serum (FCS), 2 mM L-glutamine, 100 U/ml penicillin and used to transfect one 10 cm dish with HEK 293T. After 6 h incubation the media was changed to neurobasal medium. Virus particles were harvested first 24 h after medium change, when new media was added and second after 16 h. Media contained virus was filtered using a 0.22 mm Millex GP Syringe Driven filter (Cat. n° P36359, Millipore, Billerica, MA,USA) and directly applied to neurons. Neurons were incubated with the virus at the day of plating overnight and at DIV1 either the media was changed to neurobasal medium or neurons were removed from the plating dish to be inverted on top of the astrocyte feeder layer.

### Immunocytochemistry

Neurons were fixed with 4% paraformaldehyde in PBS for 15 min and washed three times with PBS. Free aldehyde groups were quenched with 50 mM NH_4_Cl for 10 min than neurons were washed three times with PBS and permeabilized with 0.1% saponin in PBS for 15 min. Neurons were incubated in primary antibody diluted in PBS containing 0.01% saponin, 2% gelatin for 1 h. Than neurons were washed three times with PBS and incubated in secondary antibody diluted in PBS containing 0.01% saponin, 2% gelatin for 1 h. After that neurons were washed three times and mounted in microscopy slides using Mowiol.

### Live cell imaging

Neurons grown on 25 mm coverlips were used for FRET/FLIM experiments and the coverslipswere mounted in an Attofluor^®^ cell chamber for microscopy during imaging (Cat. No. A-7816, Invitrogen/molecular probes). Neurons grown on the microfluidic chambers were placed in a petri dish with an opening in the bottom and used for all motility assays. During imaging cells were placed inside a Bachhoffer heating chamber at 37°C, the pH in the medium was controlled either by adding 25 mM Hepes pH 7.4 or with 5% CO2. Fast live cell imaging of fluorescent proteins was performed using an Olympus IX71 inverted Spinning disc scan head Yokogawa CSU10. Images were acquired with a 100× /1.40 Oil UPlanSApo objective. Dual color imaging was obtained simultaneously by using a DualView image splitter HC 520/35; BS 565; HC 624/40. Images were detected by an Andor iXon EM + DU-897 BV back illuminated electron-multiplying charge-coupled device (EMCCD) image acquisition rate was from 2 to 4 frames per second.

### Image analysis of live cell imaging

The movies were imported into MotionTracking and processed as described in Rink et al ^27,60^. Briefly, to discriminate endosomes form background from cytosolic APPL1-mCherry/Akt1-GFP in axons, we applied a Bayesian foreground/background discrimination (BFBD) filter ^37^, a part of the MotionTracking software. BFBD takes advantage of slow change (high correlation) of intensities in cytoplasm over time in comparison to fast change in space of narrow axons. Movies with Akt1-GFP were processed with additional XY-bi-linear approximation due to the large amount of cytoplasmic Akt fluorescence. After BFBD filtering, the changeable component of image was discriminated (frog-eye filter). Shortly, the intensity of background estimation which was found in BFBD was subtracted from the de-noised images with uncertainty propagation to the differential image (see Supp. Movie 1 and 2). The resulting images were segmented as described in ^27,61^. Then endosomes were reliably detected by fitting the image intensity with sum of analytical functions (weighted squared Lorentzian functions). The noise was discriminated by probabilistic thresholding. Detected endosomes were associated into the tracks by dynamic programing algorithm that performs local-in-time/global-in-space optimisation ^27^. Processive movements were defined as part of track where endosome moved unidirectionally in at least 4 sequential frames. The deconvolution of histogram on log-normal components was performed by GraphAnalyzer of Motiontracking package (http://motiontracking.mpi-cbg.de). The fit was performed in range [0.07; 10] μm/sec. Optimal number of components was determined by Bayesian model probability (Occam Razor) criterion ^62^.

### Förster resonance energy transfer and fluorescence lifetime imaging

Fluorescence lifetime measurements were performed by wide-field time-domain FLIM. As an excitation light source, a Chameleon XR Ti:Sapphire Laser (Coherent) with 200 fsec pulses and a repetition rate of 90 MHz was tuned to 980 nm and frequency doubled to 490 nm using a second-harmonic generator (SHG) (APE). The SHG was equipped with an autotracker device to correct the crystal position for different wavelengths and to keep the final beam at a constant position. Finally, the laser beam was coupled into a short graded index fiber, expanded with a collimator, and coupled into the microscope. Images were recorded on a Zeiss Axiovert S100TV with a 63x 1.4 NA oil immersion objective and an optovar lens of 1.6 or 2.5 through a 495LP dichroic mirror (Chroma) and a BL HC514/30 emission band-pass filter (Semrock) for YFP fluorescence or BL HC536/40 emission band-pass filter (Semrock) for SytoxOrange emission. The images were acquired at the microscope's bottom port with an intensified PicoStar HR 12 CCD camera system (LaVision). The time-gated image intensifier was triggered directly by the laser trigger through a programmable picosecond-trigger-delay unit (LaVision). The microscope was fully automated and controlled with self-written software (LabView 8.2, National Instruments). Images were acquired and analyzed with the same software.

Fluorescence decay curves were recorded by 16 images with increasing delay times (step 500 ps) after the excitation laser pulse with a camera binning of 2 × 2 and an exposure time of 100–200 msec for each image. The laser power was reduced by ND filters to avoid photobleaching during the acquisition. Images were background corrected by subtracting the mean value of a region without any fluorescence signal, and the fluorescence lifetime was calculated for each pixel according to: ***I*_t_*=* A · e ^t/τ^** where ***I*** is the intensity measured in each pixel at a timepoint **t, A** is the preexponential factor and **τ** is the fluorescence lifetime.

### Data Availability

The datasets generated and analyzed during the current study are available from the corresponding author on reasonable request.

## Supporting information

## Author Contributions

Conceptualization, L.G.S., G.S. and M.Z.; Investigation, L.G.S., S.S. and M.P.M.; Quantitative Analysis, Y.K.; Writing, L.G.S, M.P.M and M.Z.; Writing-Reviewing and Editing, S.S. and G.S.; Funding Acquisition, M.Z.; Supervision, M.Z.

## Acknowledgements

We acknowledge Sarah Seifert and Fabian Segovia Miranda for the assistance in isolating and culturing mouse primary hippocampal neurons, Kira Bodrov for help with MotionTracking and Mike Lorenz for assistance in the FLIM imaging. Additional support came from facilities at the MPI-CBG including the light microscopy facility (LMF) and biomedical services (BMS). We thank Iain Kennedy Patten for helping write the manuscript. This work was financially supported by the Max Planck Society and LG was a recipient of a fellowship from the Christiane Nüsslein-Volhard-Foundation.

## References

1. Harrington, A. W., & Ginty, D. D. Long-distance retrograde neurotrophic factor signalling in neurons. Nat Rev Neurosci 14, 177–87 (2013).

2. Ginsberg, S. D. et al. Microarray analysis of hippocampal CA1 neurons implicates early endosomal dysfunction during Alzheimer’s disease progression. Biol Psychiatry 68, 885–93 (2010).

3. Millecamps, S. & Julien, J. P. Axonal transport deficits and neurodegenerative diseases. Nat Rev Neurosci 14, 161–76 (2013).

4. Liu, X. A., Rizzo, V. & Puthanveettil, S. V. Pathologies of axonal transport in neurodegenerative diseases. Transl Neurosci 3, 355–372 (2012).

5. Noble, W., Hanger, D. P., Miller, C. C. & Lovestone, S. The importance of tau phosphorylation for neurodegenerative diseases. Front Neurol 4, 83 (2013).

6. Wang, D., Chan, C. C., Cherry, S. & Hiesinger, P. R. Membrane trafficking in neuronal maintenance and degeneration. Cell Mol Life Sci 70, 2919–34 (2013).

7. Delcroix, J. D. et al. NGF signaling in sensory neurons: evidence that early endosomes carry NGF retrograde signals. Neuron 39, 69–84 (2003).

8. Howe, C. L. & Mobley, W. C. Signaling endosome hypothesis: A cellular mechanism for long distance communication. J Neurobiol 58, 207–16 (2004).

9. Cui, B. et al. One at a time, live tracking of NGF axonal transport using quantum dots. Proc Natl Acad Sci U A 104, 13666–71 (2007).

10. Parton, R. G., Simons, K. & Dotti, C. G. Axonal and dendritic endocytic pathways in cultured neurons. J Cell Biol 119, 123–37 (1992).

11. Schmieg, N., Menendez, G., Schiavo, G. & Terenzio, M. Signalling endosomes in axonal transport: travel updates on the molecular highway. Semin Cell Dev Biol 27, 32–43 (2014).

12. Deinhardt, K. et al. Rab5 and Rab7 control endocytic sorting along the axonal retrograde transport pathway. Neuron 52, 293–305 (2006).

13. de Renzis, S., Sönnichsen, B. & Zerial, M. Divalent Rab effectors regulate the sub-compartmental organization and sorting of early endosomes. Nat Cell Biol 4, 124–33 (2002).

14. Zerial, M. & McBride, H. Rab proteins as membrane organizers. Nat Rev Mol Cell Biol 2, 107–17 (2001).

15. Kalaidzidis, I. et al. APPL endosomes are not obligatory endocytic intermediates but act as stable cargo-sorting compartments. J Cell Biol 211, 123–44 (2015).

16. Zoncu, R. et al. A phosphoinositide switch controls the maturation and signaling properties of APPL endosomes. Cell 136, 1110–1121 (2009).

17. Zeigerer, A. et al. Functional properties of hepatocytes in vitro are correlated with cell polarity maintenance. Exp. Cell Res. 350, 242–252 (2017).

18. Murray, D. H. et al. An endosomal tether undergoes an entropic collapse to bring vesicles together. Nature 537, 107–111 (2016).

19. Wilson, J. M. et al. EEA1, a tethering protein of the early sorting endosome, shows a polarized distribution in hippocampal neurons, epithelial cells, and fibroblasts. Mol Biol Cell 11, 2657–71 (2000).

20. Luo, H. R. et al. Akt as a mediator of cell death. Proc Natl Acad Sci U A 100, 11712–7 (2003).

21. Manning, B. D. & Cantley, L. C. AKT/PKB signaling: navigating downstream. Cell 129, 1261–74 (2007).

22. Schenck, A. et al. The endosomal protein Appl1 mediates Akt substrate specificity and cell survival in vertebrate development. Cell 133, 486–497 (2008).

23. Lin, D. C. et al. APPL1 associates with TrkA and GIPC1 and is required for nerve growth factor-mediated signal transduction. Mol Cell Biol 26, 8928–41 (2006).

24. Majumdar, D., Nebhan, C. A., Hu, L., Anderson, B. & Webb, D. J. An APPL1/Akt signaling complex regulates dendritic spine and synapse formation in hippocampal neurons. Mol Cell Neurosci 46, 633–44 (2011).

25. Bonanomi, D. et al. Identification of a developmentally regulated pathway of membrane retrieval in neuronal growth cones. J Cell Sci 121, 3757–69 (2008).

26. Taylor, A. M. et al. A microfluidic culture platform for CNS axonal injury, regeneration and transport. Nat Methods 2, 599–605 (2005).

27. Rink, J., Ghigo, E., Kalaidzidis, Y. & Zerial, M. Rab conversion as a mechanism of progression from early to late endosomes. Cell 122, 735–49 (2005).

28. Villaseñor, R., Nonaka, H., Del Conte-Zerial, P., Kalaidzidis, Y. & Zerial, M. Regulation of EGFR signal transduction by analogue-to-digital conversion in endosomes. eLife 4, (2015).

29. Stenmark, H., Aasland, R., Toh, B. H. & D’Arrigo, A. Endosomal localization of the autoantigen EEA1 is mediated by a zinc-binding FYVE finger. J Biol Chem 271, 24048–54 (1996).

30. Mitsuuchi, Y. et al. Identification of a chromosome 3p14.3-21.1 gene, APPL, encoding an adaptor molecule that interacts with the oncoprotein-serine/threonine kinase AKT2. Oncogene 18, 4891–8 (1999).

31. Valdez, G. et al. Pincher-mediated macroendocytosis underlies retrograde signaling by neurotrophin receptors. J Neurosci 25, 5236–47 (2005).

32. Kaech, S. & Banker, G. Culturing hippocampal neurons. Nat. Protoc. 1, 2406–2415 (2006).

33. Miaczynska, M. et al. APPL proteins link Rab5 to nuclear signal transduction via an endosomal compartment. Cell 116, 445–56 (2004).

34. Mauro, L. et al. Evidences that estrogen receptor α interferes with adiponectin effects on breast cancer cell growth. Cell Cycle 13, 553–64 (2014).

35. Ryu, J. et al. APPL1 potentiates insulin sensitivity by facilitating the binding of IRS1/2 to the insulin receptor. Cell Rep 7, 1227–38 (2014).

36. Varsano, T. et al. GIPC is recruited by APPL to peripheral TrkA endosomes and regulates TrkA trafficking and signaling. Mol Cell Biol 26, 8942–52 (2006).

37. Morales-Navarrete, H. et al. A versatile pipeline for the multi-scale digital reconstruction and quantitative analysis of 3D tissue architecture. Elife 4, (2015).

38. Albertazzi, L., Arosio, D., Marchetti, L., Ricci, F. & Beltram, F. Quantitative FRET analysis with the EGFP-mCherry fluorescent protein pair. Photochem Photobiol 85, 287–97 (2009).

39. Macia, E. et al. Dynasore, a cell-permeable inhibitor of dynamin. Dev Cell 10, 839–50 (2006).

40. Kirchhausen, T., Macia, E. & Pelish, H. E. Use of dynasore, the small molecule inhibitor of dynamin, in the regulation of endocytosis. Methods Enzym. 438, 77–93 (2008).

41. Barroso, M. M. Quantum dots in cell biology. J Histochem Cytochem 59, 237–51 (2011).

42. Ye, M., Lehigh, K. M. & Ginty, D. D. Multivesicular bodies mediate long-range retrograde NGF-TrkA signaling. eLife 7, (2018).

43. Barford, K. et al. Transcytosis of TrkA leads to diversification of dendritic signaling endosomes. Sci. Rep. 8, 4715 (2018).

44. Schnatwinkel, C. et al. The Rab5 effector Rabankyrin-5 regulates and coordinates different endocytic mechanisms. PLoS Biol 2, E261 (2004).

45. Lorenzo, D. N. et al. A PIK3C3-ankyrin-B-dynactin pathway promotes axonal growth and multiorganelle transport. J Cell Biol 207, 735–52 (2014).

46. Naslavsky, N., Boehm, M., Backlund, P. S. & Caplan, S. Rabenosyn-5 and EHD1 interact and sequentially regulate protein recycling to the plasma membrane. Mol Biol Cell 15, 2410–22 (2004).

47. Falk, J., Konopacki, F. A., Zivraj, K. H. & Holt, C. E. Rab5 and Rab4 regulate axon elongation in the Xenopus visual system. J Neurosci 34, 373–91 (2014).

48. Deepa, S. S. & Dong, L. Q. APPL1: role in adiponectin signaling and beyond. Am J Physiol Endocrinol Metab 296, E22–36 (2009).

49. Hupalowska, A., Pyrzynska, B. & Miaczynska, M. APPL1 regulates basal NF-κB activity by stabilizing NIK. J Cell Sci 125, 4090–102 (2012).

50. Xin, X., Zhou, L., Reyes, C. M., Liu, F. & Dong, L. Q. APPL1 mediates adiponectin-stimulated p38 MAPK activation by scaffolding the TAK1-MKK3-p38 MAPK pathway. Am J Physiol Endocrinol Metab 300, E103–10 (2011).

51. Dajas-Bailador, F., Bantounas, I., Jones, E. V. & Whitmarsh, A. J. Regulation of axon growth by the JIP1-AKT axis. J. Cell Sci. 127, 230–239 (2014).

52. Cheng, H.-C. et al. AKT suppresses retrograde degeneration of dopaminergic axons by inhibition of macroautophagy. J. Neurosci. Off. J. Soc. Neurosci. 31, 2125–2135 (2011).

53. Guo, X., Snider, W. D. & Chen, B. GSK3β regulates AKT-induced central nervous system axon regeneration via an eIF2Bε-dependent, mTORC1-independent pathway. eLife 5, (2016).

54. Chaturvedi, A., Martz, R., Dorward, D., Waisberg, M. & Pierce, S. K. Endocytosed BCRs sequentially regulate MAPK and Akt signaling pathways from intracellular compartments. Nat. Immunol. 12, 1119–1126 (2011).

55. Fu, X. et al. Retrolinkin cooperates with endophilin A1 to mediate BDNF-TrkB early endocytic trafficking and signaling from early endosomes. Mol Biol Cell 22, 3684–98 (2011).

56. Del Conte-Zerial, P. et al. Membrane identity and GTPase cascades regulated by toggle and cut-out switches. Mol. Syst. Biol. 4, 206 (2008).

57. Huang, E. J. & Reichardt, L. F. Neurotrophins: roles in neuronal development and function. Annu Rev Neurosci 24, 677–736 (2001).

58. Hooper, C., Killick, R. & Lovestone, S. The GSK3 hypothesis of Alzheimer’s disease. J Neurochem 104, 1433–9 (2008).

59. Brewer, G. J. & Torricelli, J. R. Isolation and culture of adult neurons and neurospheres. Nat Protoc 2, 1490–8 (2007).

60. Chenouard, N. et al. Objective comparison of particle tracking methods. Nat Methods 11, 281–9 (2014).

61. Collinet, C. et al. Systems survey of endocytosis by multiparametric image analysis. Nature 464, 243–249 (2010).

62. Sivia, D. S. & Carlile, C. J. Molecular spectroscopy and Bayesian spectral analysis—how many lines are there? J. Chem. Phys. 96, 170 (1992).

63. Huotari J. and Helenius A., Endosome maturation, The EMBO Journal 30, 3481–3500 (2011).

64. Kim S., Sato Y., Mohan P.S., Peterhoff C., Pensalfini A., Rigoglioso A., Jiang Y, Nixon R.A., Evidence that the rab5 effector APPL1 mediates APP-βCTF-induced dysfunction of endosomes in Down syndrome and Alzheimer’s disease, Molecular Psychiatry 21, 707–716, (2016).

