## Supplementary figures and images for "Retrograde transport of Akt by a neuronal Rab5-APPL1 endosome"

### Supplementary file 1

Supp. Figure 1: APPL1 motility in hippocampal neurons

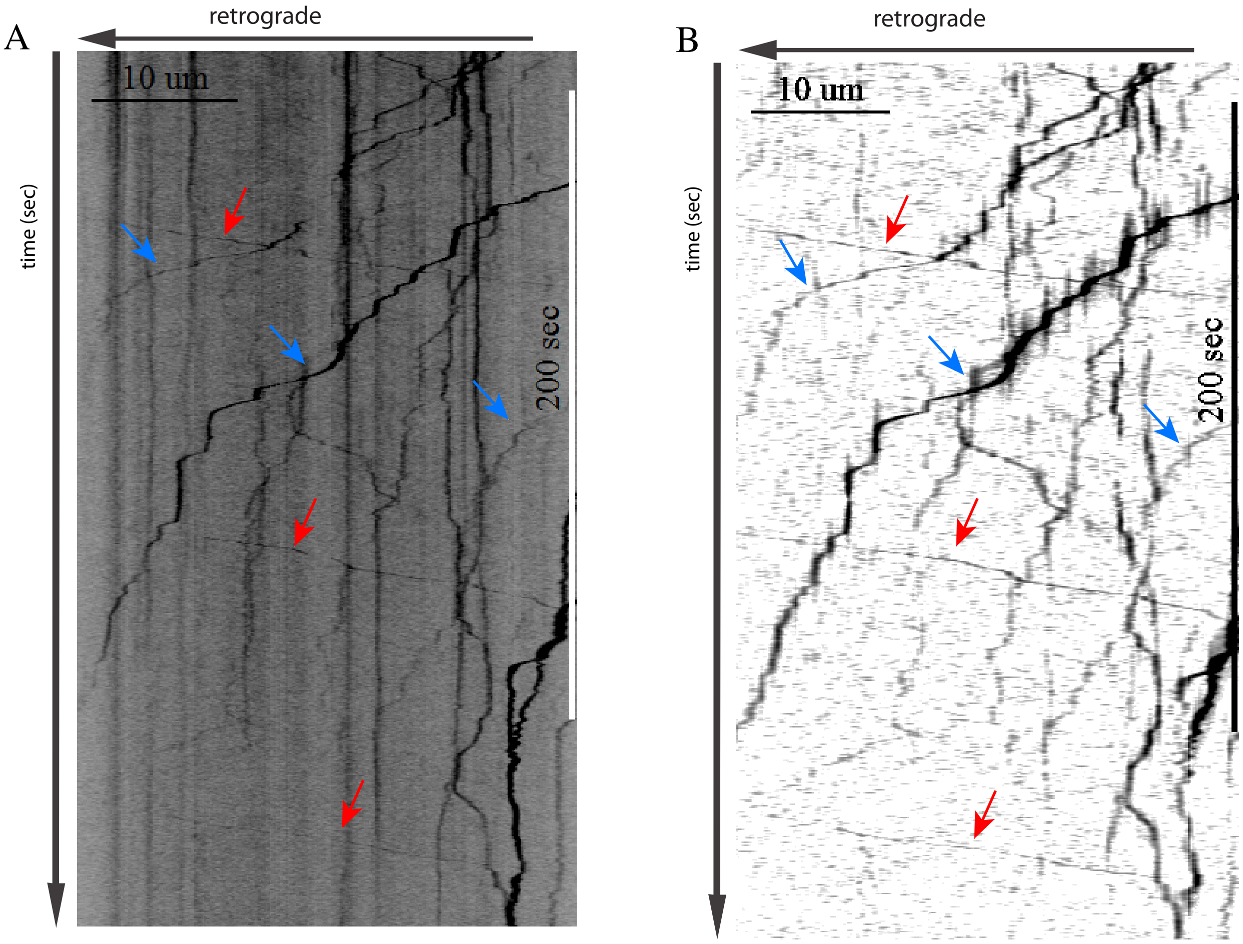

### Supplementary file 2

Supp. Figure 2: Number movement events of TrkB and APPL1 double positive endosomes

A.

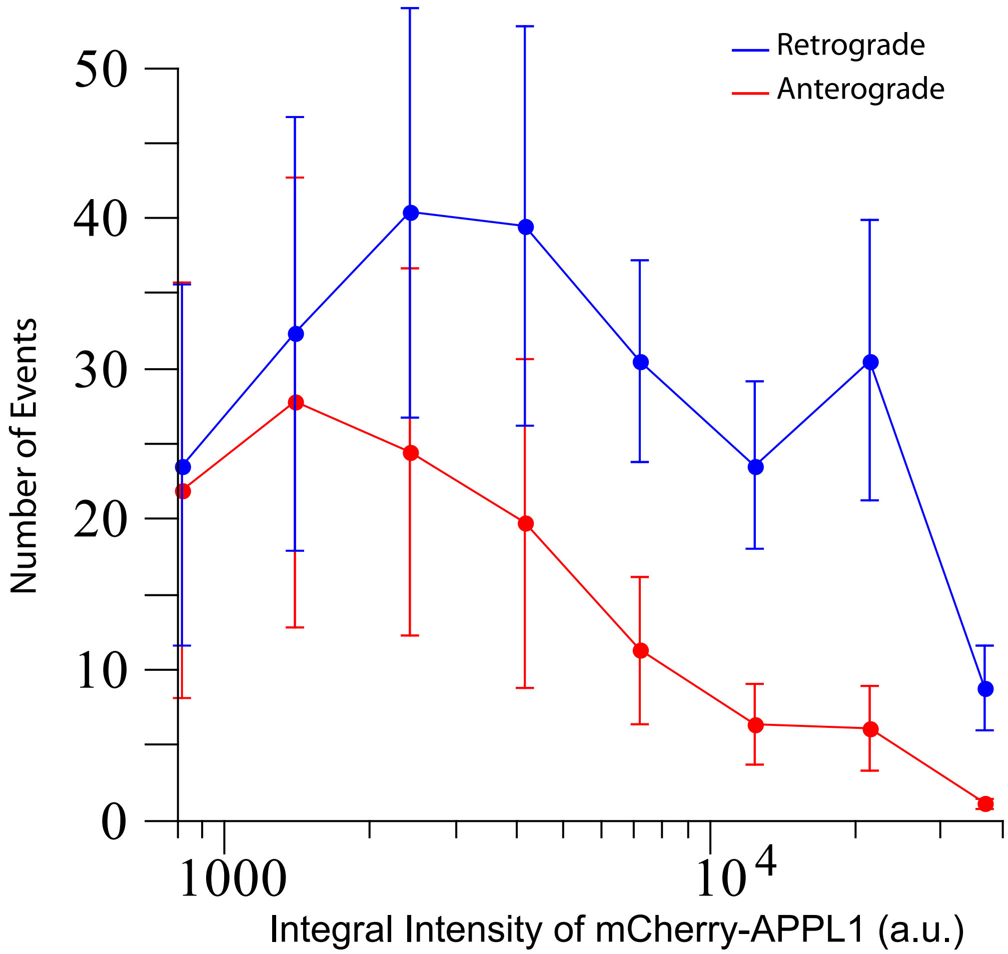
