## Supplementary material for "Retrograde transport of Akt by a neuronal Rab5-APPL1 endosome"

**Table 1: Endosomal motility data (processive tracks)**

| | Mean Size (µm) | Mean Anterograde Speed (µm/s) | Mean Retrograde Speed (µm/s) | Mean Track Displacement (µm) | Mean Processive Movement Time (s) | Mean Processive Travel Distance (µm) | Total Processive Travel Distance (µm) | # of tracks | # of cells |
|---|---|---|---|---|---|---|---|---|---|
| **Cherry-Rab5** | 0.33 [0.29 - 0.37] | 0.54 [0.48 - 0.6] | 0.87[0.63 - 1.18] | 2.5 [1.69 - 3.56] | 2.26 [1.61 - 3.09] | 1.54 [1.06 - 2.16] | 4.09 [3.52 - 4.73] | 125 | 4 |
| **Cherry-APPL1** | 0.41 [0.37 - 0.45] | 0.47 [0.32 - 0.67] | 0.48[0.37 - 0.61] | 1.46 [1.35 - 1.58] | 1.77 [1.28 - 2.38] | 0.88 [0.75 - 1.02] | 1.82 [1.70 - 1.94] | 1603 | 6 |
| **Cherry-APPL1 neurotrophic deprivation** | 0.36 [0.30 - 0.42] | 0.44 [0.30 - 0.62] | 0.54[0.40 - 0.71] | 2.02 [1.57 - 2.55] | 2.9 [2.78 - 3.02] | 1.59 [1.17 - 2.11] | 2.50 [1.87 - 3.28] | 856 | 6 |
| **Cherry-APPL1 (co-transfected with GFP-Akt)** | 0.36 [0.30 - 0.42] | 0.57 [0.43 - 0.74] | 0.72 [0.61 - 0.84] | 3.43 [2.62 - 4.42] | 3.06 [2.96 - 3.16] | 1.77 [1.43 - 2.17] | 3.41 [3.05 - 3.80] | 84 | 7 |
| **GFP-TrkB (co-transfected with mCherry-APPL1)** | 0.28 [0.26 - 0.30] | 0.46 [0.37 - 0.56] | 0.39 [0.31 - 0.48] | 1.93 [1.59 - 2.35] | 3.41 [3.09 - 3.76] | 1.28 [ 0.94 - 1.74] | 2.07 [1.68 - 2.55] | 722 | 7 |

* All values are mean [95% confidence interval]
