## Supplementary material for "Retrograde transport of Akt by a neuronal Rab5-APPL1 endosome"

**Supplementary information**

**Movie legends**

**Movie 1:** Live cell imaging of hippocampal neurons expressing mCherry-Rab5. Cell bodies localize towards the left size of the movie. Acquisition frame rate displayed in the movie. Movie played 25 frames per second.

**Movie 2:** Live cell imaging of hippocampal neurons co-expressing GFP-Rab5 and mCherry-APPL1. Acquisition frame rate displayed in the movie. Movie played 25 frames per second.

**Movie 3:** Live cell imaging of hippocampal neurons expressing mCherry-APPL1. Cell bodies localize towards the left size of the movie. Acquisition frame rate displayed in the movie. Movie played 25 frames per second.

**Movie 4:** Live cell imaging of hippocampal neurons co-expressing GFP-TrkB and mCherry-APPL1. Cell bodies localize towards the bottom of the movie. Acquisition frame rate displayed in the movie. Movie played 25 frames per second.

**Movie 5:** Live cell imaging of hippocampal neurons expressing GFP-Akt and mCherry-APPL1. Cell bodies localize towards the left size of the movie. Acquisition frame rate displayed in the movie. Movie played 25 frames per second.

**Supplementary figures and movie legends**

**Supplementary Figure 1**: Kymographs of mCherry-APPL1 endosomes in transfected primary hippocampal neurons. The neurons were grown on a microfluidic chamber. Panel A is a kymograph of movie de-noised by BFBD algorithm. Panel B presents the same kymograph after processing by FrogEye filter. FrogEye filter suppresses signal from non-moving objects and cytoplasmic background and, therefore, reveals dim moving endosomes.  The anterograde and retrograde tracks are pointed red and blue arrows respectively.

**Supplementary Figure 2**: Dependency of number of movement events in retrograde (blue) and anterograde (red) directions of double positive APPL1 and TrkB endosomes from integral intensity of mCherry-APPL1 per endosome.

**Supplementary Movie 1:** Live cell imaging of hippocampal neurons expressing mCherry-APPL1. Cell bodies localize towards the bottom of the movie. Acquisition frame rate displayed in the movie. Movie played 25 frames per second.

**Supplementary Movie 2:** The Supplementary Movie 1 after processing by FrogEye filter. Cell bodies localize towards the bottom of the movie. Acquisition frame rate displayed in the movie. Movie played 25 frames per second.
